## Supplementary tables and Supplementary figure legends for "2-deoxyglucose transiently inhibits yeast AMPK signaling and triggers glucose transporter endocytosis, potentiating the drug toxicity"

### Supplementary materials

-

### LEGENDS TO SUPPLEMENTARY TABLES.

### Table S1. Yeast strains used in this study

**Table S2.** Plasmids used in this study

**Table S3.** Antibodies used this study

### SUPPLEMENTARY TABLES.

### Table S1.

| **Name** | **Genotype & Description** | **Origin & Reference** |
| --- | --- | --- |
| ySL0066: **BY4741 (WT)** | *MAT a; ura3Δ0, his3Δ1, leu2Δ0, met15Δ0* | [86] |
| ySL0534: ***rod1Δ*** | *MAT a; ura3Δ0, his3Δ1, leu2Δ0, met15Δ0 rod1∆::KANMX* | Léon lab |
| ySL0541: ***reg1Δ*** | *MAT a; ura3Δ0, his3Δ1, leu2Δ0, met15Δ0 reg1∆::HIS3MX* | [28] |
| ySL0567: ***snf1Δ*** | *Mat alpha, ura3Δ0, his3Δ1, leu2Δ0, lys2∆0, snf1∆::kanMX4* | Léon lab |
| ySL0854: ***cnb1Δ*** | *MAT a,ura3Δ0, his3Δ1, leu2Δ0, met15Δ0 cnb1∆::KANMX* | Léon lab |
| ySL1027: **Hxt3-GFP** | *MAT a; ura3Δ0, his3Δ1, leu2Δ0, met15Δ0 HXT3::GFP-HIS3MX* | [87] |
| ySL1140: **Hxt2-GFP** | *MAT a;ura3Δ0, his3Δ1, leu2Δ0, met15Δ0 HXT2::GFP-HIS3MX* | [87] |
| ySL1186: **Hxt1-GFP** | *MAT a; ura3Δ0, his3Δ1, leu2Δ0, met15Δ0 HXT1::GFP-HIS3MX* | [87] |
| ySL1690: ***hxk2Δ*** | *MAT a; ura3Δ0, his3Δ1, leu2Δ0, met15Δ0 hxk2∆::KANMX* | Léon lab |
| ySL1852 : **Hxt4-GFP** | *MAT a; ura3Δ0, his3Δ1, leu2Δ0, met15Δ0 HXT4::GFP-HIS3MX* | [87] |
| ySL1961: ***slt2Δ*** | *MAT a; ura3Δ0, his3Δ1, leu2Δ0, met15Δ0 slt2∆::KANMX* | Euroscarf |
| ySL2092: **Vht1-GFP** | *MAT a; ura3Δ0, his3Δ1, leu2Δ0, met15Δ0 VHT1::GFP-HIS3MX* | [88] |
| ySL2110: **Pdr12-GFP** | *MAT a; ura3Δ0, his3Δ1, leu2Δ0, met15Δ0 PDR12::GFP-HIS3MX* | [88] |
| ySL2119: **Bap2-GFP** | *MAT a; ura3Δ0, his3Δ1, leu2Δ0, met15Δ0 BAP2::GFP-HIS3MX* | [88] |
| ySL2123: **Fps1-GFP** | *MAT a; ura3Δ0, his3Δ1, leu2Δ0, met15Δ0 FPS1::GFP-HIS3MX* | [88] |
| ySL2132: **Mid2-GFP** | *MAT a; ura3Δ0, his3Δ1, leu2Δ0, met15Δ0 MID2::GFP-HIS3MX* | [88] |
| ySL2142: **Qdr3-GFP** | *MAT a; ura3Δ0, his3Δ1, leu2Δ0, met15Δ0 QDR3::GFP-HIS3MX* | [88] |
| ySL2169: **Slg1-GFP** | *MAT a; ura3Δ0, his3Δ1, leu2Δ0, met15Δ0 SLG1::GFP-HIS3MX* | [88] |
| ySL2175: **Ste2-GFP** | *MAT a; ura3Δ0, his3Δ1, leu2Δ0, met15Δ0 STE2::GFP-HIS3MX* | [88] |
| ySL2204: ***hac1Δ*** | *MAT a; ura3Δ0, his3Δ1, leu2Δ0, met15Δ0 hac1∆::KANMX* | Euroscarf |
| ySL2216: **Itr1-GFP *snf1Δ*** | *MAT a; ura3Δ0, his3Δ1, leu2Δ0, met15Δ0 snf1∆::HPHNT1 Itr1::GFP-HIS3MX* | This study |
| ySL2219: **Tat1-GFP** | *MAT a; ura3Δ0, his3Δ1, leu2Δ0, met15Δ0 TAT1::GFP-HIS3MX* | [88] |
| ySL2221: **Can1-GFP** | *MAT a; ura3Δ0, his3Δ1, leu2Δ0, met15Δ0 CAN1::GFP-HIS3MX* | [88] |
| ySL2222: **Lyp1-GFP** | *MAT a; ura3Δ0, his3Δ1, leu2Δ0, met15Δ0 LYP1::GFP-HIS3MX* | [88] |
| ySL2223: **Fui1-GFP** | *MAT a; ura3Δ0, his3Δ1, leu2Δ0, met15Δ0 FUI1::GFP-HIS3MX* | [88] |
| ySL2224: **Itr1-GFP** | *MAT a; ura3Δ0, his3Δ1, leu2Δ0, met15Δ0 ITR1::GFP-HIS3MX* | [88] |
| ySL2237: **Pdr12-GFP *snf1Δ*** | *MAT a; ura3Δ0, his3Δ1, leu2Δ0, met15Δ0 PDR12::GFP-HIS3MX snf1∆::NATNT2* | This study |
| ySL2288: **Ina1-GFP** | *MAT a; ura3Δ0, his3Δ1, leu2Δ0, met15Δ0 INA1::GFP-HIS3MX* | [88] |
| ySL2315: ***hog1Δ*** | *MAT a; ura3Δ0, his3Δ1, leu2Δ0, met15Δ0 hog1∆::KANMX* | Euroscarf |
| ySL2325: **Ina1-GFP *npi1*** | *MAT a; ura3Δ0, his3Δ1, leu2Δ0, met15Δ0 INA1::GFP-HIS3MX pRSP5::KANMX* | This study |
| ySL2395: **Ina1-GFP** ***snf1Δ*** | *MAT a; ura3Δ0, his3Δ1, leu2Δ0, met15Δ0   snf1∆::HIS3MX INA1::GFP-HPHNT1* | This study |
| ySL2406: **Ina1-GFP *rod1Δ*** | *MAT a; ura3Δ0, his3Δ1, leu2Δ0, met15Δ0 INA1::GFP-HPHNT1 rod1∆::KANMX* | This study |
| ySL2412: **Ina1-GFP *rvs167Δ*** | *MAT a; ura3Δ0, his3Δ1, leu2Δ0, met15Δ0 rvs167∆KANMX4  INA1::GFP-HPHNT1* | This study |
| ySL2434: **Ina1-Δ5-GFP** | *MAT a; ura3Δ0, his3Δ1, leu2Δ0, met15Δ0 Ina1-∆5::GFP-KANMX* | This study |
| ySL2443: **Pil1-GFP** | *MAT a; ura3Δ0, his3Δ1, leu2Δ0, met15Δ0 PIL1::GFP-HIS3MX* | [88] |
| ySL2444: **Sur7-GFP** | *MAT a; ura3Δ0, his3Δ1, leu2Δ0, met15Δ0 SUR7::GFP-HIS3MX* | [88] |
| ySL2445: **Lsp1-GFP** | *MAT a; ura3Δ0, his3Δ1, leu2Δ0, met15Δ0   LSP1::GFP-HIS3MX* | [88] |
| ySL2446: **Seg1-GFP** | *MAT a; ura3Δ0, his3Δ1, leu2Δ0, met15Δ0 SEG1::GFP-HIS3MX* | [88] |
| ySL2451: **Lyp1-GFP *rod1Δ*** | *MAT a; ura3Δ0, his3Δ1, leu2Δ0, met15Δ0 LYP1::GFP-HIS3MX rod1∆::KANMX* | This study |
| ySL2452: **Qdr3-GFP *rod1Δ*** | *MAT a; ura3Δ0, his3Δ1, leu2Δ0, met15Δ0 QDR3::GFP-HIS3MX rod1∆::KANMX* | This study |
| ySL2545: **Tat1-GFP *rod1Δ*** | *MAT a; ura3Δ0, his3Δ1, leu2Δ0, met15Δ0 TAT1::GFP-HIS3MX rod1∆::KANMX* | This study |
| ySL2546: **Fui1-GFP *rod1Δ*** | *MAT a; ura3Δ0, his3Δ1, leu2Δ0, met15Δ0 FUI1::GFP-HIS3MX rod1∆::KANMX* | This study |
| ySL2580: **Vht1-GFP *rod1Δ*** | *MAT a; ura3Δ0, his3Δ1, leu2Δ0, met15Δ0 VHT1::GFP-HIS3MX rod1∆::URA3* | This study |
| ySL2591: **Ste2-GFP *rod1Δ*** | *MAT a; ura3Δ0, his3Δ1, leu2Δ0, met15Δ0 STE2::GFP-HIS3MX rod1∆::URA3* | This study |
| ySL2592: **Can1-GFP *rod1Δ*** | *MAT a; ura3Δ0, his3Δ1, leu2Δ0, met15Δ0 CAN1::GFP-HIS3MX rod1∆::URA3* | This study |
| ySL2623: **Wsc3-GFP** | *MAT a; ura3Δ0, his3Δ1, leu2Δ0, met15Δ0 WSC3::GFP-HIS3MX* | [88] |
| ySL2628: **Wsc2-GFP** | *MAT a; ura3Δ0, his3Δ1, leu2Δ0, met15Δ0 WSC2::GFP-HIS3MX* | [88] |
| ySL2671: **Ina1-GFP *hxk2Δ*** | *MAT a; ura3Δ0, his3Δ1, leu2Δ0, met15Δ0 ; INA1::GFP-HIS3MX  hxk2∆::KANMX* | This study |
| ySL2724: **Hxt1-GFP *rod1Δ*** | *MAT a; ura3Δ0, his3Δ1, leu2Δ0, met15Δ0   HXT1::GFP-HIS3MX rod1∆::HPHNT1* | This study |
| ySL2725: **Hxt3-GFP *rod1Δ*** | *MAT a; ura3Δ0, his3Δ1, leu2Δ0, met15Δ0   HXT3::GFP-HIS3MX rod1∆::HPHNT1* | This study |
| ySL2726: **Itr1-GFP *rod1Δ*** | *MAT a; ura3Δ0, his3Δ1, leu2Δ0, met15Δ0 ITR1::GFP-HIS3MX rod1∆::HPHNT1* | This study |
| ySL2727: **Pdr12-GFP *rod1Δ*** | *MAT a; ura3Δ0, his3Δ1, leu2Δ0, met15Δ0 PDR12::GFP-HIS3MX rod1∆::HPHNT1* | This study |
| ySL2728: **Lyp1-GFP *rod1Δ rog3Δ*** | *MAT a; ura3Δ0, his3Δ1, leu2Δ0, met15Δ0 LYP1::GFP-HIS3MX rod1∆::KANMX; rog3∆::NATNT2* | This study |
| ySL2739: **Vph1-mCherry** | *MAT a; ura3Δ0, his3Δ1, leu2Δ0, met15Δ0 ; Vph1-mCherry::NATNT2* | This study |
| ySL2746: ***rod1∆ hxt6∆*** | *MAT a; ura3Δ0, his3Δ1, leu2Δ0, met15Δ0 rod1∆::KANMX hxt6∆::NATNT2* | This study |
| ySL2880: **Ina1-GFP Pil1-mCherry** | *MAT a; ura3Δ0, his3Δ1, leu2Δ0, met15Δ0 INA1::GFP-HIS3MX PIL1::mCherry-NATNT2* | This study |
| ySL2965: *reg1****Δ*** | *MAT a; ura3Δ0, his3Δ1, leu2Δ0, met15Δ0 reg1∆::HPHNT2* | This study |
| ySL3003: ***rod1Δ* Hxt3-GFP** | *MAT a; ura3Δ0, his3Δ1, leu2Δ0, met15Δ0 rod1∆::KANMX Hxt3::GFP-HPHNT1* | This study |
| ySL3026: ***hxt3Δ*** | *MAT a; ura3Δ0, his3Δ1, leu2Δ0, met15Δ0 hxt3∆::KANMX* | Euroscarf |
| ySL3027: ***rod1Δhxt3Δ*** | *MAT a; ura3Δ0, his3Δ1, leu2Δ0, met15Δ0 hxt3∆::KANMX rod1∆::HPHNT1* | This study |
| ySL3028: ***hxt1Δ*** | *MAT a; ura3Δ0, his3Δ1, leu2Δ0, met15Δ0 hxt1∆::KANMX* | Euroscarf |
| ySL3029: ***rod1Δhxt1Δ*** | *MAT a; ura3Δ0, his3Δ1, leu2Δ0, met15Δ0 hxt1∆::KANMX rod1∆::HPHNT1* | This study |
| ySL3084: ***hxt1Δhxt3∆*** | *MAT a; ura3Δ0, his3Δ1, leu2Δ0, met15Δ0 hxt1∆::KANMX hxt3∆::HPHNT1* | This study |
| ySL3099: ***rod1∆ dog1∆ dog2∆*** | *MAT a; ura3Δ0, his3Δ1, leu2Δ0, met15Δ0 rod1∆::KANMX dog1∆dog2∆::LEU2* | This study |
| ySL3118 : **Hxt4-GFP *rod1∆*** | *MAT a; ura3Δ0, his3Δ1, leu2Δ0, met15Δ0 HXT4::GFP-HIS3MX rod1∆::KANMX* | This study |
| ySL3124: **Hxt2-GFP *rod1Δ*** | *MAT a; ura3Δ0, his3Δ1, leu2Δ0, met15Δ0 HXT2::GFP-HIS3MX rod1∆::HPHNT1* | This study |
| ySL3171: **Snf1-GFP** | *MAT a; ura3Δ0, his3Δ1, leu2Δ0, met15Δ0 SNF1::GFP-KANMX* | This study |
| ySL3224: ***hxk2∆*** **Snf1-GFP** | *MAT a; ura3Δ0, his3Δ1, leu2Δ0, met15Δ0 SNF1::GFP-KANMX hxk2∆::HIS3MX* | This study |
| ySL3226: ***hxk2∆*** **Hxt1-GFP** | *MAT a; ura3Δ0, his3Δ1, leu2Δ0, met15Δ0 HXT1::GFP-HIS3 hxk2∆::KANMX* | This study |
| ySL3227: ***hxk2∆*** **Hxt3-GFP** | *MAT a; ura3Δ0, his3Δ1, leu2Δ0, met15Δ0 HXT3::GFP-HIS3 hxk2∆::KANMX* | This study |
| ySL3231: ***reg1∆*** **Snf1-GFP** | *MAT a; ura3Δ0, his3Δ1, leu2Δ0, met15Δ0 SNF1::GFP-KANMX reg1∆::LEU2* | This study |

### Table S2.

| **Name** | **Description** | **Origin & Reference** |
| --- | --- | --- |
| **pSL093** | pRS416-derived (CEN, *URA3*) *p_ROD1_:ROD1*-GFP | [35] |
| **pSL094** | pRS415-derived (CEN, *LEU2*) *p_ROD1_:ROD1*-3HA | [28] |
| **pSL205** | p*_CUP1_*:6xHis-Ub (2µ, *LEU2*) (pJD421) | [89] |
| **pSL237** | pRS316-derived (CEN, *URA3*) p_Rod1_:Rod1-3Flag | Olivier Vincent |
| **pSL409** | Yep358-based (2μ, *URA3) p_DOG1_(1000bp):LacZ* | [20] |
| **pSL410** | Yep358-based (2μ, *URA3) p_DOG2_(1000bp):LacZ* | [20] |
| **pSL412** | pRS426-derived (2μ, *URA3*) *p_GPD_:DOG2* | [20] |
| **pSL436** | pRS426-derived (2μ, *URA3*) *p_GPD_:DOG2(DD>AA)* | [20] |
| **pSL559** | pRS313-derived (CEN, *HIS3*) *p_ROD1_:ROD1*-Flag | This study |
| **pSL560** | pRS313-derived (CEN, *HIS3*) *p_ROD1_:ROD1-PYm*-Flag | This study |
| **pSL561** | pRS313-derived (CEN, *HIS3*) *p_ROD1_:ROD1-S12A*-Flag | This study |
| **pSL563** | pRS313-derived (CEN, *HIS3*) *p_ROD1_:ROD1-KR*-Flag | This study |
| **pSL589** | pUG35-derived (ARS/CEN, *URA3*) *pHXT3:HXT3*-GFP | This study |
| **pSL590** | pDRf1GW-ura3-derived (2µ, *URA3*) FLII12Pglu-700μδ6 Glucose FRET sensor, Addgene #28002 | Wolf Frommer [59] |
| **pSL591** | pUG35-derived (ARS/CEN, *URA3*) *pHXT3:HXT3(N370A)*-GFP | This study |
| **pSL599** | pRS426-derived (2μ, *URA3*) *p_HXT1_*:*HXT1* | This study |
| **pSL602** | pRS426-derived (2μ, *URA3*) *p_HXT3_*:*HXT3* | This study |
| **pSL608** | pDRF1-GW-derived (2µ, *URA3*), yAT1.03 ATP FRET sensor, Addgene #132781 | Bas Teusink [49] |
| **pSL609** | pDRF1-GW-derived (2µ, *URA3*), yAT1.03 ATP FRET sensor, mutated R122K,R126K (non-ATP binding), Addgene #132782 | Bas Teusink [49] |

### Table S3.

| **Name** | **Description** | **Dilution** | **Reference/Origin** |
| --- | --- | --- | --- |
| 𝛼 GFP | Mouse monoclonal against GFP, clones 7.1/ 13.1 | 1/5000 | 11814460001 -Roche |
| 𝛼 GFP | Goat polyclonal antibody against GFP (IRDye-800 conjugated, used to reveal Snf1-GFP) | 1/2000 | 600-132-215 -Rockland |
| 𝛼 Flag | Mouse monoclonal antibody against Flag | 1/5000 | F3165 - Sigma |
| 𝛼 pAMPK / pSnf1 | Rabbit polyclonal antibody against Thr172-phosphorylated human AMPKα | 1/1000 | #2535 – Cell Signaling Technology |
| 𝛼 polyHis tag | Mouse monoclonal antibody to reveal poly-His-tagged proteins incl. Snf1 (contains a stretch of 13 His residues) | 1/2000 | H1029 - Sigma |
| 𝛼 Ubiquitin | Mouse monoclonal antibody against ubiquitin | 1/1000 | P4D1 - Santa Cruz |
| 𝛼 Rabbit IgG | Goat secondary antibody against Rabbit IgG (HRP) | 1/5000 | A6154 - Sigma |
| 𝛼 Mouse IgG | Goat secondary antibody against Mouse IgG (HRP) | 1/5000 | A5278 - Sigma |

### LEGENDS TO SUPPLEMENTARY FIGURES

**Figure S1 - 2DG treatment induces vacuolar fragmentation**. **A.** WT cells expressing Vph1 tagged with mCherry were grown overnight in glucose medium (exponential phase) and treated with 0.2% 2DG. Cells were collected and observed by fluorescence microscopy at the indicated times. Scale bar: 5 µm. **B.** Quantification of 2DG-induced vacuolar fragmentation (values ± SD, *n=*3 independent experiments).

**Figure S2 - Most of the studied plasma membrane proteins are endocytosed in response to 2DG and this endocytosis is often Rod1-dependent.** WT or *rod1∆* cells expressing the indicated membrane proteins tagged with GFP at their endogenous genomic locus were observed by fluorescence microscopy before and after treatment with 2DG for 4h. Scale bar, 5 µm.

**Figure S3 - Rog3 is not responsible for the observed Rod1-independent endocytosis of Lyp1-GFP in response to 2DG.** WT, *rod1∆* and *rod1∆ rog3∆* cells expressing Lyp1-GFP tagged at its endogenous genomic locus were observed by fluorescence microscopy before and after treatment with 2DG for 4h. Scale bar, 5 µm.

**Figure S4- Expression of Rod1-3HA does not restore endocytosis nor sensitivity to 2DG in a *rod1∆* context, contrary to Rod1-Flag**. (**A**) The indicated strains were observed by fluorescence microscopy after growth in a glucose-containing medium and after 2DG treatment for 4h. Scale bar, 5 µm. (**B**) Serial dilutions of cultures of the indicated mutants/plasmid combinations were spotted on the indicated media and grown for 3 days at 30°C.

**x**

**Figure S5- Ina1-GFP is endocytosed in response to 2DG.** (**A**) WT or *rvs167∆* strains expressing Ina1-GFP were observed by fluorescence microscopy after growth in a glucose-containing medium and after 2DG treatment for 4h. In addition, WT cells were incubated with CMAC to label the vacuole. Scale bar, 5 µm. (**B**) *Left*, Total protein extracts of WT cells expressing Ina1-GFP were prepared before and after 2h or 4h 2DG treatment and immunoblotted using anti-GFP antibodies. *Right*, the same extracts were treated with endoglycosidase H and immunoblotted using anti-GFP antibodies.

**Figure S6- Endocytosis of Hxt1-GFP and Hxt3-GFP requires Rod1 interaction with Rsp5 and Rod1 ubiquitylation.** The indicated strains were observed by fluorescence microscopy after growth in a glucose-containing medium and after 2DG treatment for 4h. Scale bar, 5 µm.

**Figure S7- *SNF1* deletion is not sufficient to block endocytosis in response to 2DG**. Pdr12-GFP, Pdr12-GFP *snf1∆*, Itr1-GFP and Itr1-GFP *snf1∆* cells were grown in a glucose-containing medium and observed by fluorescence microscopy before and after 2DG treatment for 4h. Scale bar, 5 µm

**Figure S8- Snf1-GFP as a tool to follow Snf1 phosphorylation and function. A.** Serial dilutions of cultures of the WT, *reg1∆*, *snf1∆* and Snf1-GFP strains were spotted on synthetic complete medium containing either glucose or sucrose as a carbon source and grown for 3 days at 30°C. **B.** Total protein extracts of WT, Snf1-GFP, *reg1*∆, *reg1*∆ Snf1-GFP and *snf1∆* cells expressing Mig1-Flag were prepared after overnight growth (exponential phase) in glucose medium (H: high glucose), after transfer to 0.05% glucose for 2h (L: low glucose) and after addition of glucose for 10 min (+Glc). Samples were immunoblotted using anti-Flag, anti-GFP and anti-pAMPK antibodies.MW: molecular weight marker.

**Figure S9- *HXK2* deletion or *DOG2* overexpression prevent Hxt1 and Hxt3 endocytosis in response to 2DG**. **A.** Hxt1-GFP, Hxt1-GFP *hxk2∆*, Hxt3-GFP and Hxt3-GFP *hxk2∆*cells were grown in a glucose-containing medium and observed by fluorescence microscopy before and after 2DG treatment for 4h. Scale bar, 5 µm. **B.** Strains expressing Hxt1-GFP and Hxt3-GFP containing an empty plasmid (Ø) or plasmids allowing the overexpression of *DOG2* or its catalytic mutant *DOG2-DDAA* were grown in a glucose-containing medium and observed by fluorescence microscopy before and after 2DG treatment for 4h. Scale bar, 5 µm.
